# PickET: An unsupervised method for localizing macromolecules in cryo-electron tomograms

**DOI:** 10.1101/2025.08.20.671250

**Authors:** Shreyas Arvindekar, Omkar Golatkar, Shruthi Viswanath

**Affiliations:** National Center for Biological Sciences, Tata Institute of Fundamental Research, Bangalore, India 560065

**Keywords:** Cryo-electron tomography, unsupervised learning, particle-picking, localization, protein complexes, macromolecular assemblies

## Abstract

Cryo-electron tomography (cryo-ET) datasets are rich sources of information capable of describing the localizations, structures, and interactions of macromolecules. However, most current methods for localizing particles in cryo-electron tomograms are limited to macromolecules with known structure, require extensive manual annotations, and/or are computationally expensive. Here, we present PickET, a method for localizing macromolecules in tomograms that does not rely on expert annotations and prior structures. Its performance is demonstrated on a diverse dataset comprising over a hundred tomograms from publicly available datasets, varying in sample types, sample preparation conditions, microscope hardware, and image processing workflows. We demonstrate that PickET can simultaneously localize macromolecules of various shapes, sizes, and abundance. The predicted particle localizations can be used for 3D classification and *de novo* structural characterization. Our fully unsupervised approach is efficient and scalable, and enables high-throughput analysis of cryo-ET data.

## Introduction

Cryo-electron tomography (cryo-ET) is a cryo-EM imaging modality that enables the visualization of entire cells or lamellae milled from them at nanometer resolutions (Ng & Gan, 2020). In contrast with the traditionally used reductionist approach in structural biology, cryo-ET enables the structural characterization of macromolecules in their native cellular environment (Asano et al., 2016; Beck & Baumeister, 2016; Gubins et al., 2020; Lamm et al., 2022; Ng & Gan, 2020). The generated tomograms are rich sources of information as they provide a spatial description of the cellular proteome at nanometer and potentially sub-nanometer resolution (Mahamid et al., 2016; Xue et al., 2022). Comprehensively mapping macromolecules in tomograms to characterize their spatial distributions and interactions to build molecular atlases of cells is known as visual proteomics (Asano et al., 2016; Beck & Baumeister, 2016; Förster et al., 2010; Robinson et al., 2007; Turk & Baumeister, 2020). A major bottleneck in visual proteomics lies in localizing macromolecules in tomograms (Arvindekar et al., 2024; Majila et al., 2025). The missing wedge, the low signal-to-noise ratio in the data, the crowded cellular environment, and the considerable variation among particles in terms of size, shape, and abundance in tomograms make this task challenging (Y. Chen et al., 2014; Gubins et al., 2020; Lučić et al., 2013; Maurer et al., 2024b; Moebel et al., 2021; Pyle & Zanetti, 2021).

Template matching (TM) is a commonly used approach for localizing macromolecules with known structures in tomograms (Böhm et al., 2000; Frangakis et al., 2002; Maurer et al., 2024b; Pyle & Zanetti, 2021). In TM, a low-pass filtered reference template of a macromolecule is used to obtain the positions and orientations of the target macromolecule in the tomogram (Böhm et al., 2000; Frangakis et al., 2002; Maurer et al., 2024b). Several software packages have been developed for template matching, including recent ones that show significant improvements in accuracy and efficiency (Castaño-Díez et al., 2012, 2017; Cruz-León et al., 2024; Hrabe et al., 2012; Lucas et al., 2021; Maurer et al., 2024a, 2024b; Pyle & Zanetti, 2021; Wan et al., 2024). However, TM is limited to macromolecules with known structures, does not account for conformational and compositional heterogeneity, and often fails to distinguish between structurally similar particles (Jin et al., 2024; Kim et al., 2023; Maurer et al., 2024b). In addition, localizing small particles using TM, especially in crowded environments, is still challenging (Cruz-León et al., 2024; Martinez-Sanchez, 2025). Moreover, the high false positive rate often requires complex and time-consuming post-processing (Cruz-León et al., 2024). Lastly, TM is computationally expensive and has low throughput, limiting its usability for visual proteomics (Martinez-Sanchez, 2025).

To overcome the limitations of TM, several deep learning-based approaches have been developed (M. Chen et al., 2017; de Teresa-Trueba et al., 2023; Huang et al., 2024; Liu et al., 2024; Moebel et al., 2021; Rice et al., 2023; Uddin et al., 2024; Zeng et al., 2018, 2023). Supervised learning-based approaches have been shown to outperform TM (M. Chen et al., 2017; de Teresa-Trueba et al., 2023; Gubins et al., 2020; Lamm et al., 2022, 2024; Last et al., 2024; Moebel et al., 2021; Wagner et al., 2019). However, these methods often do not generalize across domains (*e.g.*, different specimen types and microscope hardware). Usually, they can localize only the macromolecules used in their training, and often fail to localize less abundant particles (Cruz-León et al., 2024). Moreover, they require a considerable amount of reliable training data, which is time-consuming, labor- and compute-intensive to obtain, since it relies on template matching and/or manual annotation (Martinez-Sanchez, 2025).

To extend the usability of deep learning-based methods by reducing the dependence on extensive annotations, several supervised and self-supervised representation learning-based methods have been developed (Huang et al., 2024; Rice et al., 2023; Zeng et al., 2018, 2023). In these methods, features describing subvolumes from the input tomogram are learnt by a neural network. These features are then used to cluster subvolumes containing structurally similar particles. Further, fully unsupervised methods have also been developed (Jin et al., 2024; Martinez-Sanchez et al., 2020; Uddin et al., 2025). In (Martinez-Sanchez et al., 2020; Uddin et al., 2025), the features for clustering subvolumes are not learnt, but instead derived from discrete Morse theory and pre-trained models from computer vision, respectively. Although unsupervised methods hold great promise for enabling high-throughput particle localization in tomograms, their use in routine particle localization is still limited, as they rely on substantial manual input and require improvements in both efficiency and accuracy (Cruz-León et al., 2024; Huang et al., 2024; Martinez-Sanchez, 2025; Wagner & Raunser, 2025).

Here, we present PickET, a library of workflows for high-throughput unsupervised localization of macromolecules in cryo-electron tomograms that does not rely on extensive training data or prior structures of macromolecules. It is efficient, scalable to large datasets, and requires minimal user input. We demonstrate the performance of PickET on 133 tomograms from five publicly available datasets that vary in the biological specimens studied, sample types imaged, sample thinning process used, imaging hardware used, image processing software employed for generating the reconstructions, and the particle types annotated (de Teresa-Trueba et al., 2023; Dietrich et al., 2022; Ermel et al., 2024; Harrington et al., 2024; Khavnekar et al., 2023; Peck et al., 2024; Rice et al., 2023). PickET can simultaneously localize macromolecules of various shapes, sizes, and abundance. It is comparable to another self-supervised particle localization method, MiLoPYP (Huang et al., 2024), outperforming the latter on a recent dataset generated using the advanced plasma-FIB milling technology. PickET predictions can be used for 3D classification using methods such as ReLiON (Scheres, 2012) and *de novo* structural characterization, as well as to supplement template matching and manual annotations for developing advanced particle localization methods. Efficient and scalable unsupervised methods for localizing macromolecules in tomograms, such as PickET, can contribute to high-throughput visual proteomics studies.

## Results

### Overview of PickET

PickET is a modular Python library for unsupervised localization of particles in cryo-electron tomograms. A typical PickET run on an input tomogram involves two steps (**Fig. 1, Materials and Methods**). The first step generates a semantic segmentation that separates the voxels associated with particles from the ones associated with the background in the input tomogram. Features describing each voxel are extracted from a 3D sub-volume or neighborhood around the voxel. These features are then used to cluster the voxels into two groups—particles and background—resulting in a binary semantic segmentation. In the second step, the binary semantic segmentation is converted to an instance segmentation, distinguishing individual particle instances. The geometric centroids of the particle instances in the instance segmentations are the predicted particle localizations. Owing to the modular architecture of the PickET library, each of the three feature extraction modes – Intensities, Fast Fourier Transform (FFT), and Gabor – can be paired with either of the clustering algorithms – K-Means and Gaussian Mixture Models (GMM) – providing six semantic segmentations for the input tomogram in the first step. The six semantic segmentations can be processed with either of the two particle extraction algorithms – connected component labeling (CC) and watershed segmentation (WS) – in the second step, providing twelve sets of particle localizations corresponding to the twelve workflows in PickET (See **Materials and Methods**).

**Figure 1:**
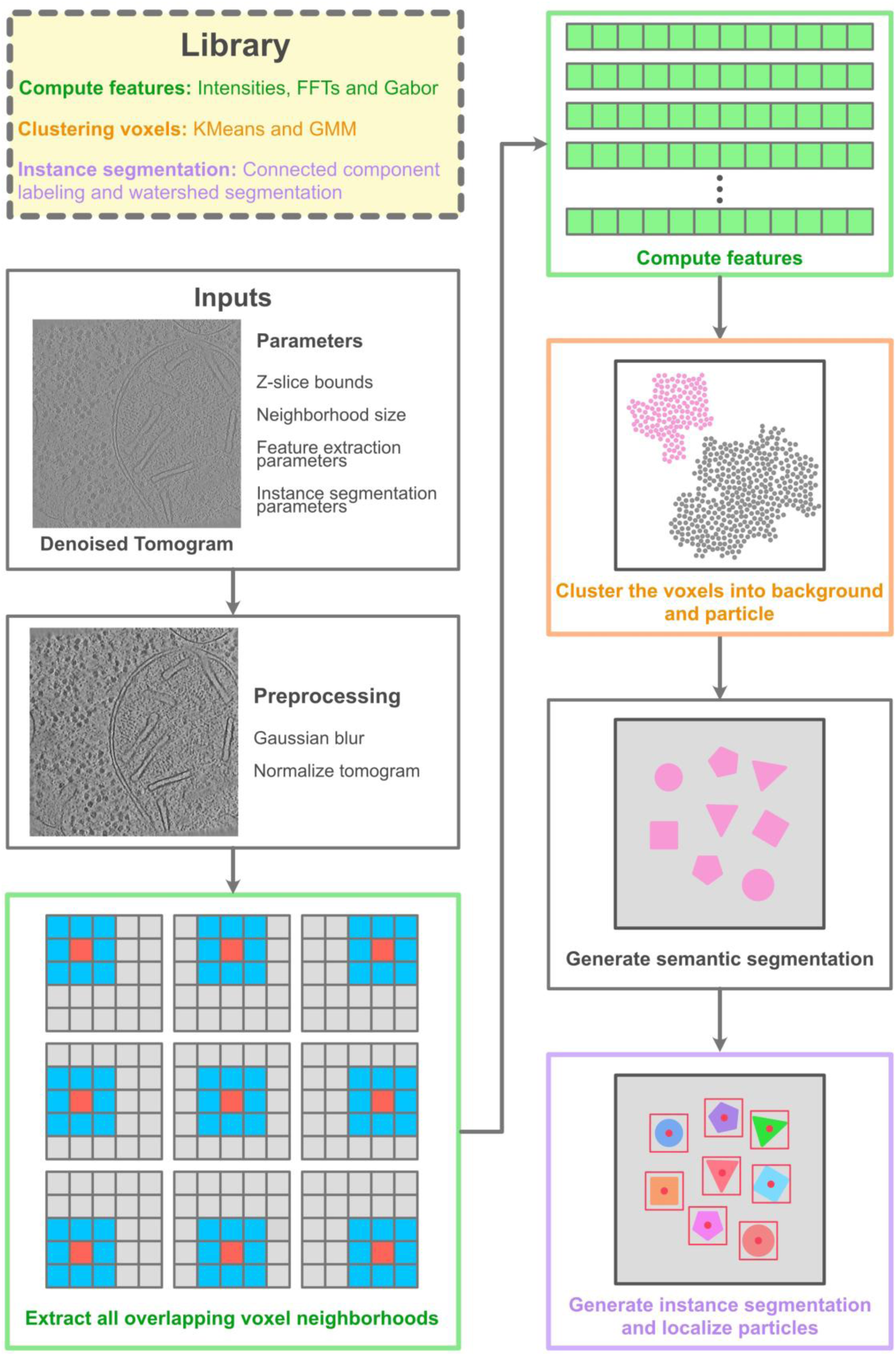
PickET library. Given an input tomogram, PickET uses voxel neighborhood-based workflows to localize particles in the tomogram in an unsupervised manner. Each PickET workflow consists of two steps, with the feature extraction (green boxes) and clustering (orange box) modules comprising the first (generating semantic segmentation) step, and the particle extraction module (purple box) comprising the second (localizing particles) step.

### PickET localizes macromolecules in tomograms

First, we compared the PickET workflows on a dataset comprising 88 simulated tomograms in terms of their precision, recall, and F1-score (**Fig. 2A**, **Table 1**, **Fig. S1, Materials and Methods, Assessment**). On the simulated tomogram dataset, all the PickET workflows performed better than the random baseline (**Fig. 2A, Fig. S1**). Among the workflows, Gabor-GMM-CC (and Intensities-GMM-CC) performed the best with median F1-scores of 0.72 (and 0.71), respectively (**Fig. 2A, Fig. S1, Fig. S2**). These workflows were also efficient, taking an average of ∼12 minutes and ∼26 minutes per tomogram, respectively (**Fig. S3A**). We show an example of semantic and instance segmentation using Gabor-GMM-CC on the simulated dataset (**Fig. 2B, 2C**).

**Figure 2:**
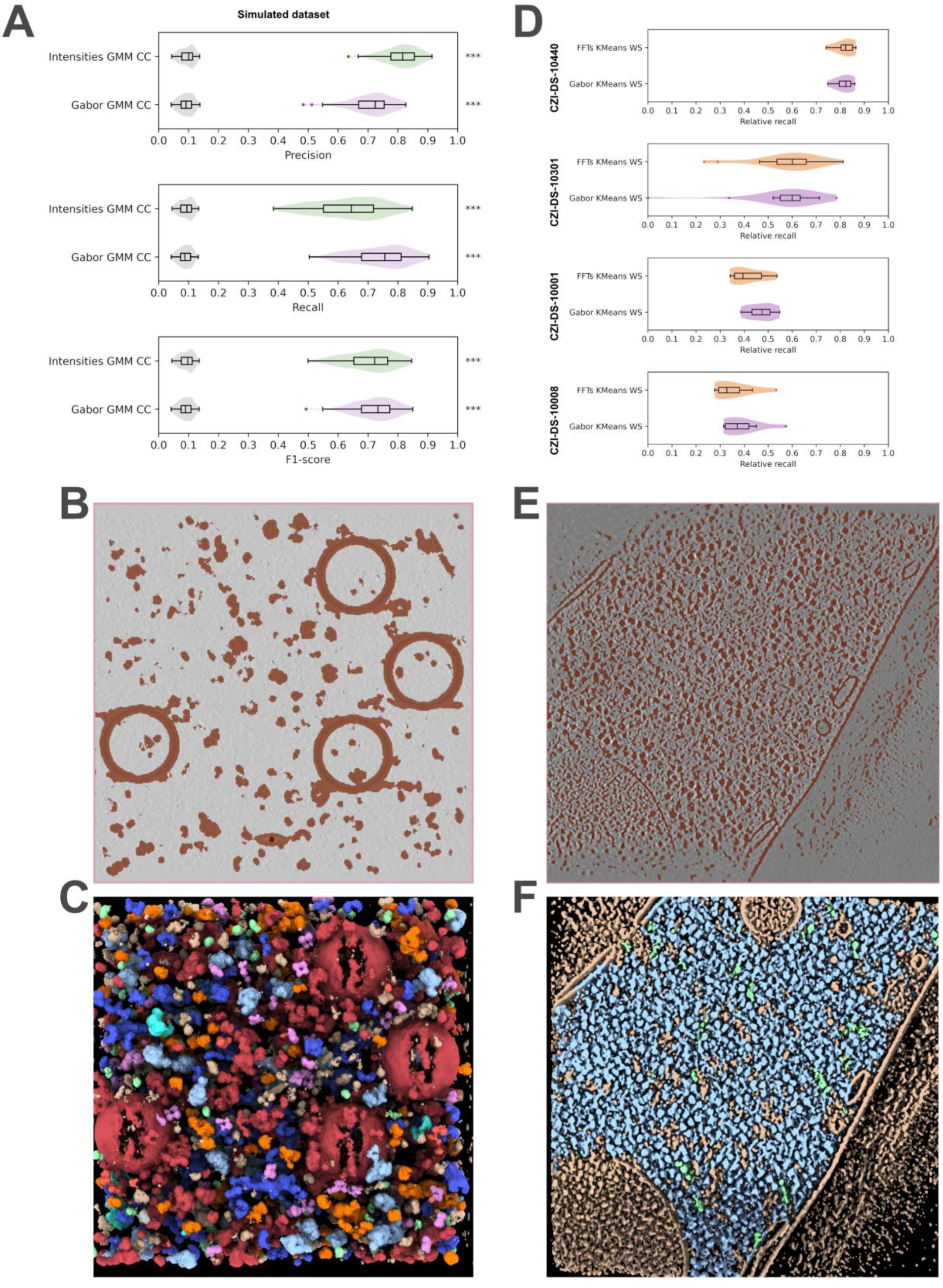
Particle localization performance of PickET. The performance of the best-performing PickET workflows (green, purple, and orange violins) on a dataset comprising simulated tomograms (*n* = 88) (**A**) and four real-world datasets (CZI-DS-10440 (*n* = 7), CZI-DS-10301 (*n* = 18), CZI-DS-10001 (*n* = 10), and CZI-DS-10008 (*n* = 9)) (**D**) is shown. On simulated tomograms, the performance of PickET was also compared against a random baseline (gray violins) (**A**). Central Z-slices of the binary semantic segmentation in brown from a PickET workflow for a simulated tomogram (**B**) and a real-world tomogram from CZI-DS-10001 (**E**). The segmentations from PickET are shown on the corresponding simulated and real-world tomograms (**C, F**). Particles are color-annotated according to the ground truth annotations. Cream-colored segmentations represent particles annotated by PickET and missing from the ground truth. Note that (**C** and **F**) are 3D visualizations of the particle segmentation, whereas (**B** and **E**) are 2D visualizations of the semantic segmentation on the central Z-slice. The statistical significance of the difference between the metrics calculated on the model predictions and random predictions is represented by asterisks (‘n.s.’ - *p* − *value* > 0.05, ‘*’ - 0.05 ≥ *p* − *value* > 0.01, ‘**’ - 0.01 ≥ *p* − *value* > 0.001, and ‘***’ - 0.001 ≥ *p* − *value*).

**Table 1:**
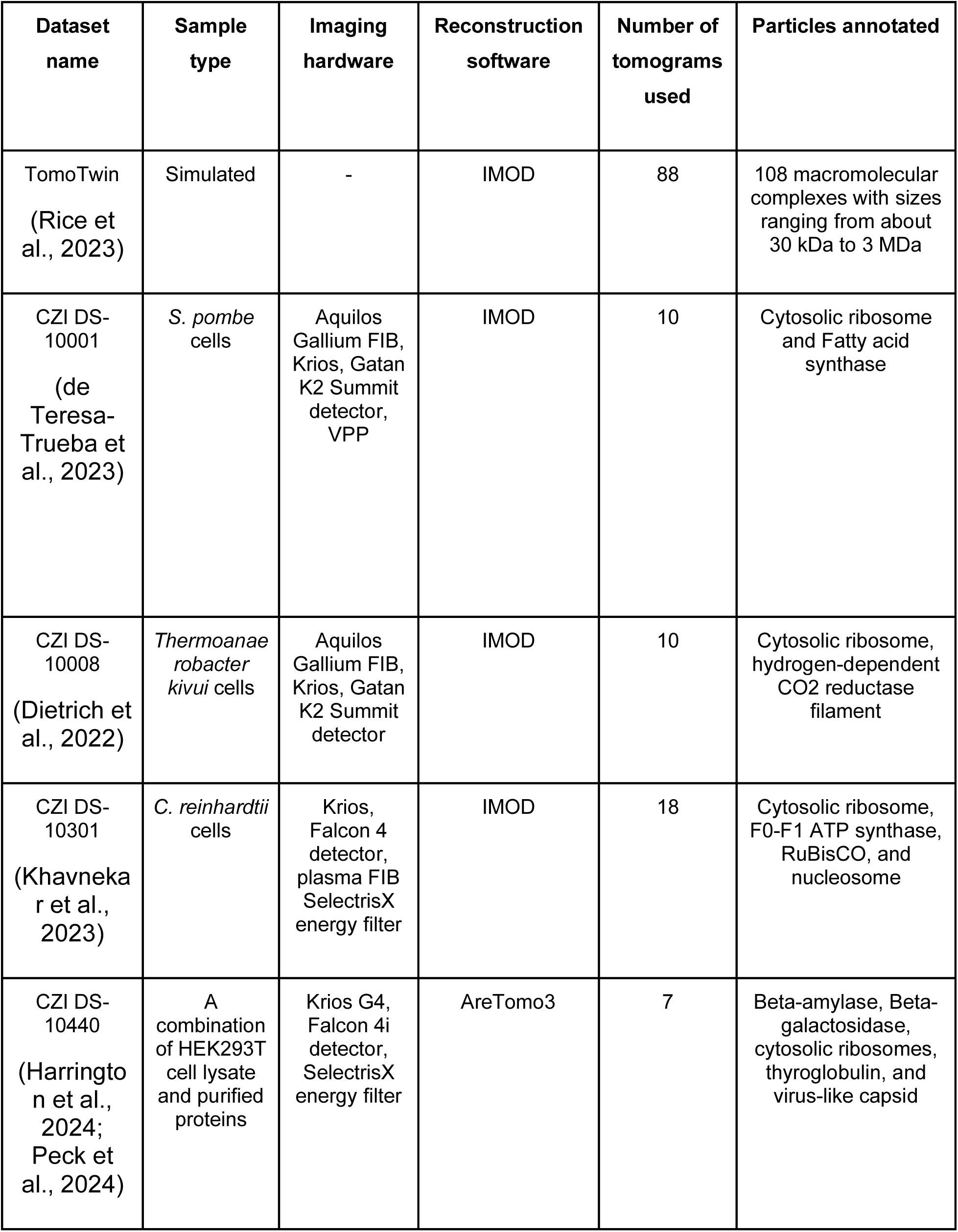
Datasets used for evaluating PickET and competing methods.

Next, we compared the PickET workflows on a dataset comprising 45 real-world tomograms (**Table 1, Materials and Methods, Datasets**). For real-world datasets, the ground truth annotations were available only for a few particle types. Whereas, PickET is not limited to specific particle types. With incomplete annotations, the precision may be underestimated, leading to a misleading F1-score. As an alternative, we used relative recall that balances the recall on the PickET predictions with that of random guessing for the same number of predictions, without explicitly relying on precision (**Materials and Methods, Assessment**). Among the workflows, Gabor-KMeans-WS (and FFT-KMeans-WS) performed the best, with median relative recalls of about 0.82 (0.82), 0.57 (0.58), 0.47 (0.42), and 0.39 (0.36) for the lysate (CZI-DS-10440), plasma-FIB milled *C. reinhardtii* (CZI-DS-10301), Gallium-FIB-milled *S. pombe* (CZI-DS-10001), and Gallium-FIB-milled *T. kivui* (CZI-DS-10008) datasets, respectively **(Fig. 2D, Fig. S4, Fig. S5, Table 1).** Among the best-performing workflows, Gabor-KMeans-WS was the most efficient, requiring, on average, approximately 15 minutes per tomogram on the CZI-DS-10301 dataset (**Fig. S3B**). We show example semantic and instance segmentations using Gabor-KMeans-WS on real-world datasets (**Fig. 2E, 2F, Fig. 3**).

**Figure 3:**
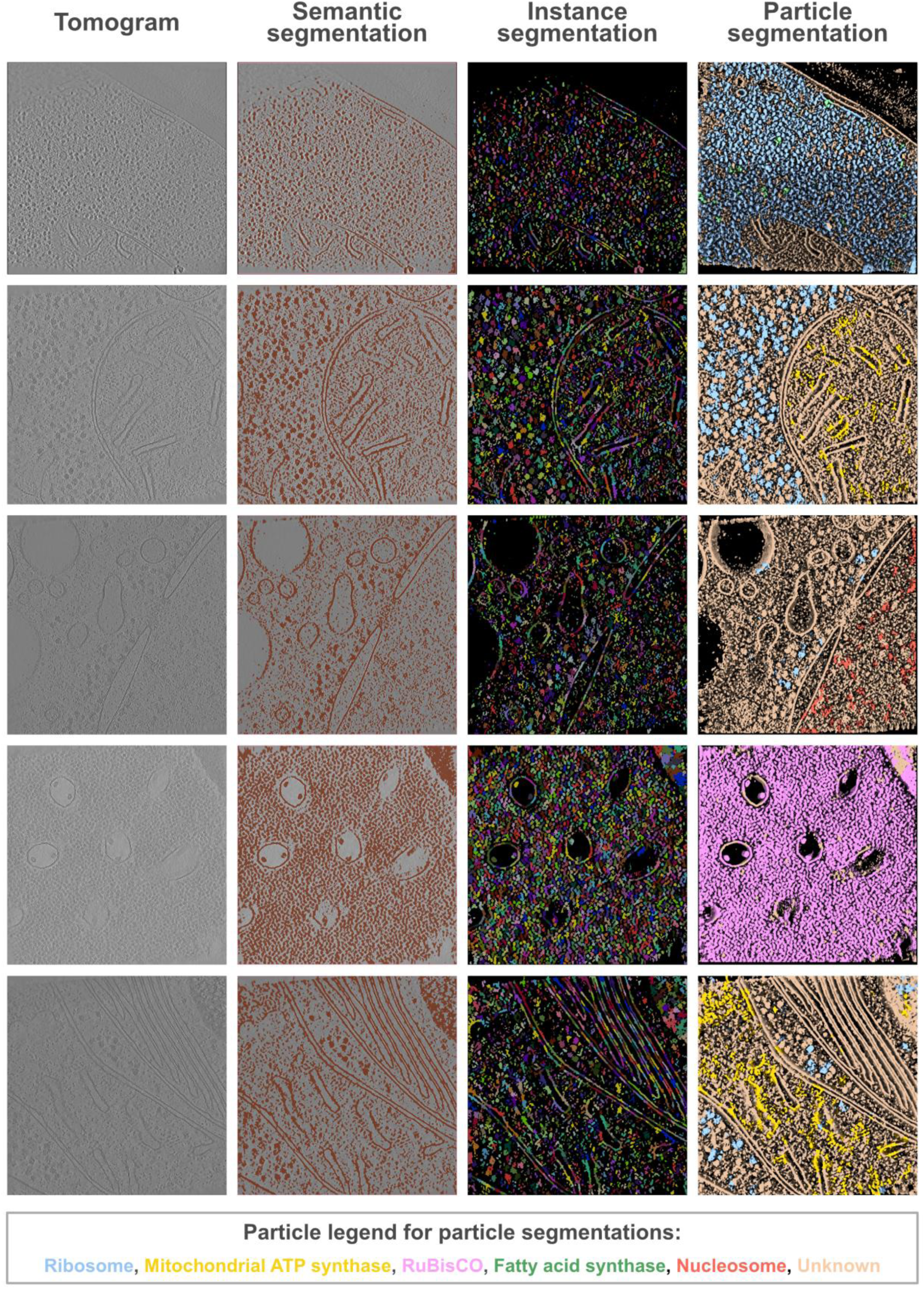
Representative segmentations from PickET. The first, second, and third columns show a Z-slice through the input tomogram, the semantic segmentation overlaid on the tomogram, and the instance segmentation – from the Gabor-KMeans-Watershed workflow. The fourth column is a 3D visualization of the instance segmentation around the Z-slice shown in the previous columns. Segmentations that overlapped with a ground truth annotation are colored by the corresponding ground truth particle type; segmentations without a corresponding ground truth annotation are marked as “Unknown” (see legend). The tomograms are from CZI-DS-10001 (first row: gallium FIB milled *S. pombe* lamella, with ground truth annotations for cytosolic ribosomes and fatty acid synthase) and CZI-DS-10301 (subsequent rows: plasma FIB milled *C. reinhardtii* lamellae, with ground truth annotations for cytosolic ribosomes, nucleosomes, RuBisCO, and mitochondrial ATP synthase). Note that the last column shows 3D visualizations of the particle segmentation, whereas the second and third columns are 2D visualizations of the segmentation on the central Z-slice.

PickET can localize large, abundant particles with known structures, such as ribosomes, that can also be annotated by template matching (**Fig. 3, Fig. S6, Table S1**). In addition, it can also localize small, less abundant particles (**Fig. S6, Table S1**). Further, the instance segmentations suggest that PickET also localizes particles that were not annotated in the ground truth, indicating that it can be used to supplement the annotations (**Fig. 2C, 2F, Fig. 3**). In summary, PickET can simultaneously and efficiently localize macromolecules of various sizes, shapes, and abundance (**Fig. 3**, **Fig. S6, Table S1**).

### Comparison to another particle localization method

Next, we compared the best-performing PickET workflows – Gabor-GMM-CC for simulated tomograms and Gabor-KMeans-WS for the real-world tomograms – with MiLoPYP, another particle localization method (**Fig. 4**). For this comparison, only the first step (Cellular Content Exploration) of MiLoPYP was used (**Materials and Methods, Comparison with existing methods**). The two methods were compared on all 133 tomograms used in this study based on the precision, recall, and F1-score on the simulated dataset and the relative recall on the real-world datasets (**Table 1**, **Fig. 4**). In addition, the two methods were compared based on the number of particles predicted and the total time taken (**Fig. 4**).

**Figure 4:**
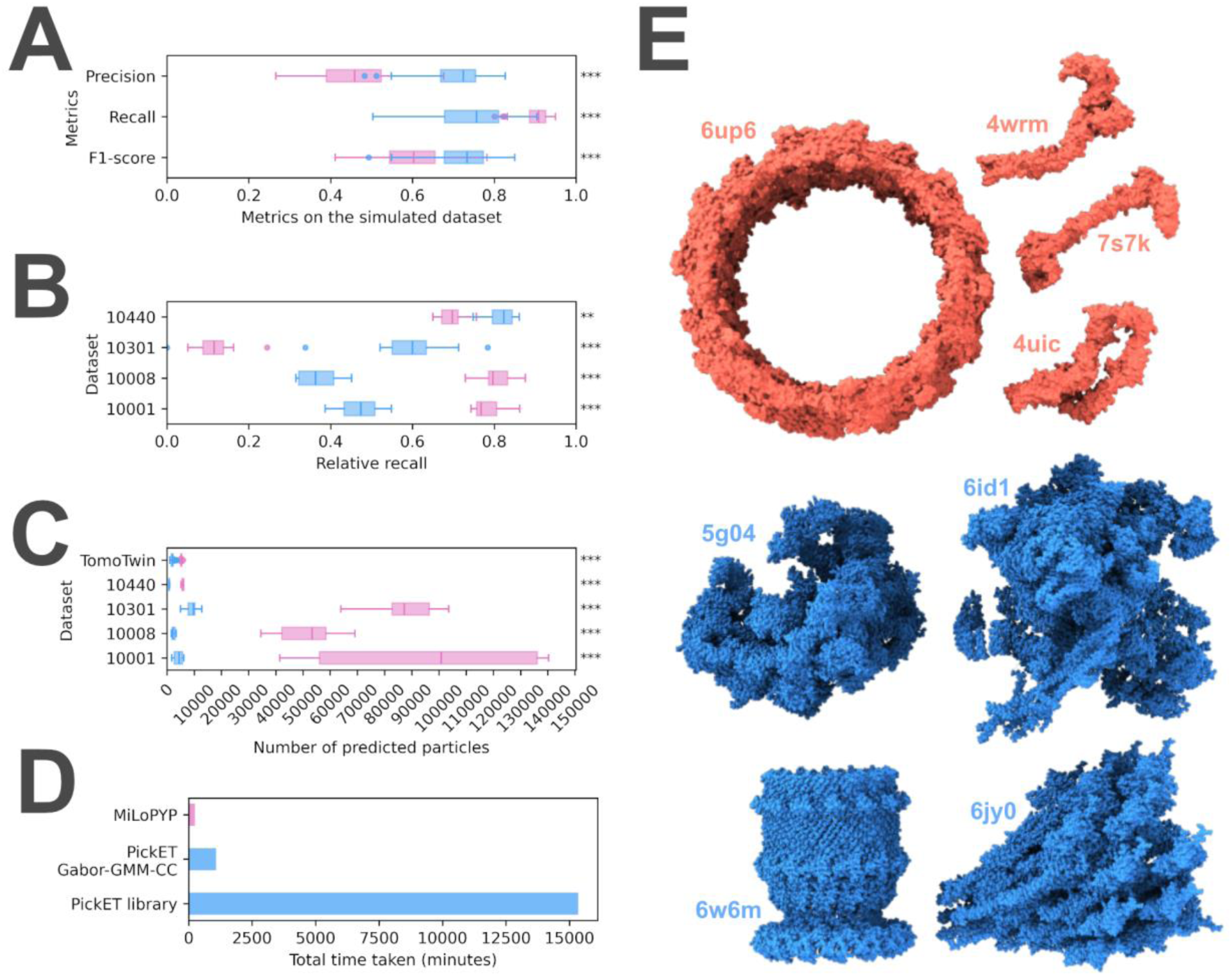
Comparison between the performance of PickET and MiLoPYP. PickET (blue) and MiLoPYP (pink) were compared based on their precision, recall and F1-score on simulated tomograms (*n* = 88) **(A)**; relative recall on the real world tomograms (*n* = (7,18,10,9) for CZI-DS-10440, CZI-DS-10301, CZI-DS-10001, and CZI-DS-10008, respectively) **(B)**; the number of particles predicted on all the tomograms **(C)**; and their run times **(D).** The runtime comparison was on 88 simulated 512 ∗ 512 ∗ 200 voxel tomograms, measured on an AMD Ryzen Threadripper 3990X workstation CPU (64 core, 2.9 GHz, 256 GB RAM) with NVIDIA RTX Ada 6000 GPU. **(E)** The particles on which both methods performed poorly (well) are shown in red (blue). The statistical significance of the difference between the metrics calculated on the predictions from the two methods is represented by asterisks (‘n.s.’ - *p* − *value* > 0.05, ‘*’ - 0.05 ≥ *p* − *value* > 0.01, ‘**’ - 0.01 ≥ *p* − *value* > 0.001, and ‘***’ - 0.001 ≥ *p* − *value*).

On the simulated dataset, PickET had a higher precision, lower recall, and higher F1 score compared to MiLoPYP (**Fig. 4A**). On the lysate and the plasma-FIB-milled *C. reinhardtii* datasets (CZI-DS-10440 and CZI-DS-10301, respectively), PickET performed significantly better than MiLoPYP in terms of the relative recall. On the remaining two gallium-FIB-milled *S. pombe* and *T. kivui* datasets (CZI-DS-10001 and CZI-DS-10008, respectively), MiLoPYP performed considerably better than the selected PickET workflow (**Fig. 4B**). Overall, MiLoPYP predicted significantly more particles than PickET, which resulted in a higher recall and lower precision in its predictions (**Fig. 4A-4C**). In terms of efficiency, the full PickET library as well as the single selected workflow take more time than MiLoPYP (**Fig. 4D**). This may be attributed to the tomogram-specific design of PickET, which processes one tomogram at a time. The selected PickET workflow (Gabor-KMeans-WS) took ∼1080 minutes versus ∼244 minutes for MiLoPYP on 88 simulated tomograms on a workstation (**Fig. 4D**). However, the efficiency of PickET can be easily improved in at least two ways. One, by trivially parallelizing runs across tomograms and across workflows. Two, by using a small subset of tomograms to estimate the clustering parameters before applying them to the entire dataset. In summary, PickET performs comparably to MiLoPYP, with MiLoPYP performing better on tomograms from gallium FIB-milled samples. In contrast, PickET performs better on lysates and, importantly, it significantly outperforms MiLoPYP on the tomograms from the more advanced plasma FIB-milled workflows.

### Influence of particle characteristics on localization performance

First, we examined how the performance of PickET varies with particle size. About 77% (83 out of 108) of the macromolecules in the simulated dataset, ranging in size from 60 kDa to 2.8 MDa, were localized by PickET with an average particle-wise recall of at least 0.7. Out of 62 macromolecules with molecular weight greater than 500 kDa, 50 (∼81%) were localized by PickET with an average particle-wise recall of at least 0.7 (**Fig. S6A**), showing that PickET is reasonably accurate at localizing medium to large macromolecules. Moreover, out of 46 macromolecules smaller than 500 kDa, 33 (∼72%) were localized by PickET with an average particle-wise recall of at least 0.7, demonstrating that PickET can also localize small macromolecules reasonably well. A similar trend was seen for the dependence of particle-wise recall on radius of gyrations (**Fig. S6B**). Similarly, we also demonstrated the particle-wise recall for PickET in the real-world datasets (**Table S1**). In summary, PickET can localize macromolecules of various sizes and shapes.

Second, we sought to identify which particle shapes are easier or more challenging to pick. Interestingly, both PickET and MiLoPYP showed poor recall on the same particle types (**Fig. 4E**). For both methods, particles that were small, hollow and/or had an elongated (4wrm, 7s7k, 4uic), or donut-like (6up6) shape were challenging to localize; whereas, large, compact particles (5g04, 6id1, 6w6m, 6jy0) were localized well by both methods (**Fig. 4E**).

## Discussion

Here, we developed PickET, a library of workflows for fully unsupervised localization of macromolecules in cryo-electron tomograms. It can be used in conjunction with 3D classification and subtomogram averaging for the *de novo* structural characterization of macromolecules. Given the efficiency and precision of PickET (**Fig. 2A, Fig. S3**), it can be used to supplement template matching and manual annotations for training deep learning-based particle localization and identification methods. In addition, PickET-generated semantic segmentations can be used in area-selective template matching approaches (Last et al., 2024) to improve the localization of target macromolecules in tomograms.

The tomograms generated from different sample types, sample thinning methods, and microscope hardware often vary in their signal-to-noise ratio, making it challenging to develop a particle-picking method that generalizes well across datasets. PickET bypasses this issue with its tomogram-specific design. We demonstrated the generalizability of PickET on 133 tomograms that vary in terms of particle types annotated, biological specimens studied, sample types being imaged, sample thinning process, imaging hardware used, and image processing software employed for generating the reconstructions (**Table 1**). PickET can simultaneously localize macromolecules of various shapes and sizes, ranging from 60 kDa to 2.8 MDa; even localizing several macromolecules smaller than 500 kDa (**Fig. S6, Table S1**). Importantly, PickET can localize macromolecules irrespective of their abundance (**Fig. 3, Table S1**). Moreover, PickET does not require known structures. Additionally, it is efficient, scalable to large datasets, and can be applied in a high-throughput manner. Therefore, owing to its generalizability, its non-reliance on previously characterized structures, and its ability to localize macromolecules of various shapes, sizes, and abundance, PickET can serve as a valuable tool for visual proteomics studies.

The performance on the tomograms from real-world datasets provided a realistic view of PickET’s applicability. The real-world tomograms were more complex compared to the simulated dataset. The latter contained a sparse distribution of particles and virtually no contaminants, such as ice crystals. On the contrary, the real-world tomograms illustrated the common challenges with tomograms, such as the crowded intracellular landscape, and artifacts from sample preparation, such as uneven lamellae and the presence of contaminants. The comparatively worse performance of the workflows from the PickET library on some of the real-world tomograms may be attributed to these challenges.

For crowded tomograms used in this study (such as CZI-DS-10001, CZI-DS-10008, and CZI-DS-10301), the Gabor-KMeans-WS workflow performed the best, whereas for comparatively sparse tomograms (such as from the simulated dataset and CZI-DS-10440), Gabor-GMM-CC worked the best (**Fig. S1-S5**). Across all datasets, FFTs-GMM-CC performed the worst, predicting considerably more particles than other workflows (**Fig. S1-S5**). This could be attributed to the dusty semantic segmentations that it generates, wherein small groups of background voxels are falsely segmented as particles. For generating semantic segmentations on crowded tomograms, K-Means clustering performed better than GMM, as the latter often resulted in larger particle segmentations that merged segmentations of neighboring particles. Similarly, for generating instance segmentations on crowded tomograms, watershed segmentation-based workflows were better suited than connected component labeling, as the latter often failed to separate neighboring particles.

In general, PickET generates good semantic segmentations that effectively separate the particle- and background-associated voxels (**Fig. 2**, **Fig. 3**). However, the methods used for generating instance segmentations often perform poorly in crowded environments. Additionally, the centroid-based particle localizations used in PickET may not be appropriate for localizing elongated, branched, or lobed particles (**Fig. 4**). This highlights the need for better instance segmentation and particle annotation methods.

An assumption in the unsupervised semantic segmentation step of PickET is that there is a background-particle separation. However, the clustering algorithm may instead separate fiducials, contaminants, or membranes from the rest of the tomogram. Moreover, artifacts such as unevenness in the lamellae and contaminants such as ice crystals make it challenging to distinguish particles from the background. PickET includes the option to crop the tomogram along the Z-axis before semantic segmentation, to reduce the effect of artifacts on the clustering, as well as before instance segmentation, to limit the localization to the lamella. In the future, additional preprocessing steps may be added to PickET to remove the voxels corresponding to such damages before generating the semantic segmentations. Finally, the voxel clustering-based approach used in PickET is memory-intensive. To address this, we provide options such as setting the Z-slice bounds and subsampling voxels before generating semantic segmentations in PickET to make it adaptable to the available memory.

Recent advancements in hardware and software for imaging cells using cryo-ET have led to an increase in the quality and volume of tomography data (Martinez-Sanchez, 2025). High-throughput particle localization methods that do not rely on ground truth annotations, such as PickET, hold great promise for advancing visual proteomics. The datasets made available by the Cryo-ET Data Portal (https://cryoetdataportal.czscience.com/) (Ermel et al., 2024) will further accelerate the development and evaluation of such methods.

## Materials and Methods

### Datasets

To evaluate the performance of PickET, we compiled a dataset by combining the tomograms and particle annotations from several publicly available datasets. Overall, the compiled dataset comprised 133 tomograms, including simulated and real-world cases (**Table 1**). The annotations on these tomograms encompass over a hundred different types of particles, ranging in size from about 30 kDa to over 3 MDa. The real-world tomograms in the dataset were carefully chosen to ensure diversity in terms of particle types annotated, biological specimens studied (bacteria, yeasts, mammalian cells), sample types being imaged (lysates, whole cells, and cellular lamellae), specimen processing (milling process used for obtaining the lamellae – *e.g.*, gallium or plasma FIB), imaging hardware used, and image processing software employed for generating the reconstructions.

### PickET workflow

PickET is a modular library for unsupervised localization of particles in a cryo-electron tomogram, comprising three feature extraction modes, two clustering algorithms, and two particle extraction algorithms. We define a PickET workflow as a specific selection of a feature extraction mode, a clustering algorithm, and a particle extraction algorithm. Any feature extraction mode can be used along with either of the clustering algorithms and either of the particle extraction algorithms, providing a total of twelve workflows. Each such workflow is split into two steps, with the feature extraction and clustering modules comprising the first (generating semantic segmentation) step, and the particle extraction module comprising the second (localizing particles) step (**Fig. 1**).

#### Generating semantic segmentation

The first step in all the workflows in the PickET library is to generate binary semantic segmentations for the input tomogram that classify each voxel as belonging to a particle or background.

##### Inputs

The input tomogram is first denoised using TomoEED with default settings (https://sites.google.com/site/3demimageprocessing/tomoeed, (Moreno et al., 2018)). Often, tomograms feature a central slab along the Z-axis, where the tomogram is the most informative, *i.e.*, it is most likely to contain particles. We provide an option to specify the bounds on this central Z-slab to generate semantic segmentations. Using these bounds reduces the memory requirement for larger tomograms and may also help avoid the contaminants from the periphery of the lamella from confounding the clustering algorithm. Additional options are provided for reducing the memory requirements for larger tomograms (See https://github.com/isblab/pickET).

The semantic segmentation generation step can be further divided into three sections: preprocessing, feature extraction, and clustering.

##### Preprocessing

The input tomogram is first blurred using a Gaussian kernel with a standard deviation *σ* = 2 to enhance contrast. Then, the voxel intensities of the blurred tomogram are normalized to be between zero and one.

##### Feature extraction

Owing to the low signal-to-noise ratio in cryo-electron tomograms, classifying the voxels based solely on their intensity values often results in segmentations that fail to distinguish particles from the background. We found that a sub-volume around each voxel, termed the neighborhood of a voxel, provides more informative context for characterizing a voxel. We use *N*_*s*_ × *N*_*s*_ × *N*_*s*_ = 5 × 5 × 5 voxel neighborhoods around all non-peripheral voxels to compute their features; a non-peripheral voxel is at least two voxels (⌊*N*_*s*_ /2⌋ = 2) away from the edge of the tomogram. Out of the three feature extraction modes provided in the PickET library – intensities, FFT, and Gabor – the last two are GPU-accelerated.

##### Intensities mode

In the intensities mode, the intensities corresponding to all 125 voxels in the 5 × 5 × 5 neighborhood of a voxel are used as its features.

##### FFT mode

In the FFT mode, the magnitudes of the 3D discrete Fourier transform of each voxel neighborhood are used as the features of a voxel.

##### Gabor mode

In the Gabor mode, first, a Gabor filter bank containing *m* Gabor filters is generated. A Gabor filter is a 3D matrix *G ∈ R*^*Ns*×*Ns*×*Ns*^, where *N*_*s*_ = 5 is the size of the voxel neighborhood. It is generated as the product of a 3D Gaussian distribution and a sinusoid. An element *G*(*x*, *y*, *z*) in such a matrix is computed as

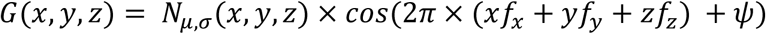

Where, *x*, *y*, and *z* represent indices in the filter along the *X*, *Y*, and *Z* axes, *N*_*μ*,*σ*_ is a Gaussian distribution with mean *μ* and standard deviation *σ*, and *f*_*x*_, *f*_*y*_, and *f*_*z*_ are the frequencies of the sinusoid along the respective axes. In the Gabor filters used in PickET, μ, the mean of the Gaussian is placed at the centre of the filter, *i.e*., at (⌊*N*_*s*_/2⌋, ⌊*N*_*s*_/2⌋, ⌊*N*_*s*_/2⌋) = (2,2,2), and σ, the standard deviation, is set to be (⌊*N*_*s*_/2⌋/ 3 = 2/3. The phase *ψ* is set to 0. For this study, ten sinusoidal frequencies were used per axis, spaced uniformly between 0 and the Nyquist frequency (0.5 cycles/voxel), resulting in a filter bank comprising *m* = 10 × 10 × 10 = 1000 filters. Convolving the filter bank with voxel neighborhoods yields *m* responses per voxel. We then select a subset, *m*_*out*_ = 64 filters with the highest response standard deviations across all voxels to define the Gabor features for a voxel.

##### Clustering

Based on the features, the voxels are clustered into two clusters using either of the two clustering algorithms – K-Means or GMM. Usually, the two clusters correspond to background and particle classes. By default, the smaller (larger) cluster is considered to represent the voxels associated with particles (background). However, given that this assumption may not hold for a small subset of segmentations, PickET includes an option to invert the generated cluster assignments.

Clustering may be performed in two steps if the tomogram is too large to be processed in one step or if central Z-slab bounds are provided (see Inputs). In such cases, first, the cluster parameters are estimated based on the central Z-slab. Then, these parameters are used to assign clusters to all voxels, resulting in a binary semantic segmentation, with particle (background) voxels marked with 1 (0).

#### Localizing particles

The second step in all the workflows in the PickET library is to localize particles in the tomogram, given a binary semantic segmentation as input. Based on the input semantic segmentation, instance segmentations that separate individual particle instances are generated using one of two particle extraction algorithms – connected component labeling (CC) and watershed segmentation (WS). Geometric centroids of individual particle instances are then computed as the predicted particle localizations.

As a result, a run of the complete PickET library generates six semantic segmentations in the first step, and twelve instance segmentations with corresponding particle localizations in the second step for each input tomogram. An optimal instance segmentation is one in which, first, the particles are well separated from the background and, second, the individual particle instances are well separated from each other. The corresponding particle localizations may be used for downstream tasks such as subtomogram averaging.

### Assessment

#### Precision, recall, and F1-score

PickET workflows were assessed using precision, recall, and F1-score on the simulated dataset. For the simulated dataset, a predicted particle localization was considered a true positive if the centroid of the predicted particle was within 100 Å of a ground truth particle centroid. Similarly, this distance threshold was set to 125 Å for predictions on the real-world datasets to account for the error in the particle localizations in the ground truth information.

#### Relative recall

PickET workflows are not limited to specific particle types. However, the ground truth annotations in real-world datasets were available only for a few particle types. This implied that the ground truth annotations for assessing the performance of methods like PickET on real-world datasets were incomplete. In cases with incomplete ground truth, the absence of a ground truth annotation in the vicinity of a prediction suggests one of three possibilities: ground truth annotations were not generated on the particle type captured by PickET, ground truth annotations were generated for this particle type, but this instance was missed by the annotation method, or the PickET prediction is a false positive. In such cases, the precision may be underestimated, leading to a potentially misleading F1-score.

The relative recall is an alternative metric to assess the performance of PickET workflows on real-world datasets with incomplete ground truth annotations, without relying on precision. Relative recall balances the recall on the PickET predictions with the recall for the same number of predictions made by random guessing.

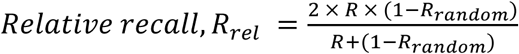

Where *R*_*rel*_ is the relative recall, *R* is the recall for PickET predictions, and *R*_*random*_ is the recall for the same number of particle localizations predicted at random. Relative recall ranges from 0 to 1, with values close to 1 indicating that PickET performs better than random guessing, and values near 0 indicating no improvement over random.

A comparison between the relative recall and F1-score on a simulated setup revealed that for a constant precision, the F1-score increased with an increase in the number of predictions. Whereas, relative recall decreased with an increase in the number of predictions, penalizing the model for overpredicting relative to the ground truth annotations **(Fig. S7)**.

### Comparison with existing methods

The performance of PickET was compared with that of MiLoPYP (dataset-specific cellular pattern **Mi**ning and particle **Lo**calization **PY**thon **P**ipeline), a self-supervised learning-based method on 125 tomograms (**Table 1)** (Huang et al., 2024). MiLoPYP is a two-step method. The first step, cellular content exploration, uses self-supervised contrastive learning for localizing particles. The second step, protein-specific particle localization, uses few-shot learning to localize particles of a specific type (for instance, ribosomes), based on a subset of user-picked particles from the first step. Since the second step is particle-specific, we compared our method using only the output from the first step of MiLoPYP. The best-performing PickET workflows, as identified earlier (**Fig. 2**, **Fig. 3, Fig. S1-S5**), were compared to MiLoPYP based on the precision, recall, and F1-score for the simulated tomograms and relative recall for the real-world tomograms (**Fig. 4**).

## Acknowledgements

We thank Kartik Majila and Muskaan Jindal for their useful comments on the manuscript. We are thankful to Thorsten Wagner and Markus Stabrin for their valuable suggestions on the development of the method. In addition, we thank Kartik Majila for the suggestions on the assessment of the method. We are also grateful to the Cryo-ET Data Portal (https://cryoetdataportal.czscience.com/) (Ermel et al., 2024), built by the Chan Zuckerberg Imaging Institute and the Chan Zuckerberg Initiative, for providing the real-world cryo-ET datasets used in this study. Molecular graphics images were produced using Napari (Sofroniew et al., 2025) UCSF ChimeraX packages from the Resource for Biocomputing, Visualization, and Informatics at the University of California, San Francisco (supported by NIH P41 RR001081, NIH R01-GM129325, and National Institute of Allergy and Infectious Diseases).

## Conflict of interest

None declared.

## Funding

This work was supported by the following grants: Department of Atomic Energy (DAE) TIFR grant RTI 4006 and Department of Biotechnology (DBT) grant BT/PR40323/BTIS/137/78/2023 from the Government of India to S.V.

## Data availability

All the simulated tomograms used in this study were from (Rice et al., 2023). The real-world tomograms were obtained from the Cryo-ET Data Portal (https://cryoetdataportal.czscience.com/) with the dataset IDs mentioned in Table 1. The segmentations and corresponding input tomograms shown in the figures are available at https://doi.org/10.5281/zenodo.16909581.

## Code availability

The Python-based source code and a detailed documentation for PickET are available at https://github.com/isblab/pickET.

## Notes

https://github.com/isblab/pickET

### Competing Interest Statement

The authors have declared no competing interest.

## References

Arvindekar, S., Majila, K., & Viswanath, S. (2024). Recent methods from statistical inference and machine learning to improve integrative modeling of macromolecular assemblies (Version 4). arXiv. 10.48550/ARXIV.2401.17894

Asano, S., Engel, B. D., & Baumeister, W. (2016). *In Situ* Cryo-Electron Tomography: A Post-Reductionist Approach to Structural Biology. Journal of Molecular Biology, 428(2, Part A), 332–343. 10.1016/j.jmb.2015.09.030

Beck, M., & Baumeister, W. (2016). Cryo-Electron Tomography: Can it Reveal the Molecular Sociology of Cells in Atomic Detail? Trends in Cell Biology, 26(11), 825–837. 10.1016/j.tcb.2016.08.006

Böhm, J., Frangakis, A. S., Hegerl, R., Nickell, S., Typke, D., & Baumeister, W. (2000). Toward detecting and identifying macromolecules in a cellular context: Template matching applied to electron tomograms. Proceedings of the National Academy of Sciences, 97(26), 14245–14250. 10.1073/pnas.230282097

Castaño-Díez, D., Kudryashev, M., Arheit, M., & Stahlberg, H. (2012). *Dynamo*: A flexible, user-friendly development tool for subtomogram averaging of cryo-EM data in high-performance computing environments. Journal of Structural Biology, 178(2), 139–151. 10.1016/j.jsb.2011.12.017

Castaño-Díez, D., Kudryashev, M., & Stahlberg, H. (2017). *Dynamo Catalogue*: Geometrical tools and data management for particle picking in subtomogram averaging of cryo-electron tomograms. Journal of Structural Biology, 197(2), 135–144. 10.1016/j.jsb.2016.06.005

Chen, M., Dai, W., Sun, S. Y., Jonasch, D., He, C. Y., Schmid, M. F., Chiu, W., & Ludtke, S. J. (2017). Convolutional neural networks for automated annotation of cellular cryo-electron tomograms. Nature Methods, 14(10), 983–985. 10.1038/nmeth.4405

Chen, Y., Pfeffer, S., Fernández, J. J., Sorzano, C. O. S., & Förster, F. (2014). Autofocused 3D Classification of Cryoelectron Subtomograms. Structure, 22(10), 1528–1537. 10.1016/j.str.2014.08.007

Cruz-León, S., Majtner, T., Hoffmann, P. C., Kreysing, J. P., Kehl, S., Tuijtel, M. W., Schaefer, S. L., Geißler, K., Beck, M., Turoňová, B., & Hummer, G. (2024). High-confidence 3D template matching for cryo-electron tomography. Nature Communications, 15(1), 3992. 10.1038/s41467-024-47839-8

de Teresa-Trueba, I., Goetz, S. K., Mattausch, A., Stojanovska, F., Zimmerli, C. E., Toro-Nahuelpan, M., Cheng, D. W. C., Tollervey, F., Pape, C., Beck, M., Diz-Muñoz, A., Kreshuk, A., Mahamid, J., & Zaugg, J. B. (2023). Convolutional networks for supervised mining of molecular patterns within cellular context. Nature Methods, 20(2), Article 2. 10.1038/s41592-022-01746-2

Dietrich, H. M., Righetto, R. D., Kumar, A., Wietrzynski, W., Trischler, R., Schuller, S. K., Wagner, J., Schwarz, F. M., Engel, B. D., Müller, V., & Schuller, J. M. (2022). Membrane-anchored HDCR nanowires drive hydrogen-powered CO2 fixation. Nature, 607(7920), 823–830. 10.1038/s41586-022-04971-z

Ermel, U., Cheng, A., Ni, J. X., Gadling, J., Venkatakrishnan, M., Evans, K., Asuncion, J., Sweet, A., Pourroy, J., Wang, Z. S., Khandwala, K., Nelson, B., McCarthy, D., Wang, E. M., Agarwal, R., & Carragher, B. (2024). A data portal for providing standardized annotations for cryo-electron tomography. Nature Methods, 21(12), 2200–2202. 10.1038/s41592-024-02477-2

Förster, F., Han, B.-G., & Beck, M. (2010). Chapter Eleven—Visual Proteomics. In G. J. Jensen (Ed.), Methods in Enzymology (Vol. 483, pp. 215–243). Academic Press. 10.1016/S0076-6879(10)83011-3

Frangakis, A. S., Böhm, J., Förster, F., Nickell, S., Nicastro, D., Typke, D., Hegerl, R., & Baumeister, W. (2002). Identification of macromolecular complexes in cryoelectron tomograms of phantom cells. Proceedings of the National Academy of Sciences, 99(22), 14153–14158. 10.1073/pnas.172520299

Gubins, I., Chaillet, M. L., van der Schot, G., Veltkamp, R. C., Förster, F., Hao, Y., Wan, X., Cui, X., Zhang, F., Moebel, E., Wang, X., Kihara, D., Zeng, X., Xu, M., Nguyen, N. P., White, T., & Bunyak, F. (2020). SHREC 2020: Classification in cryo-electron tomograms. Computers & Graphics, 91, 279–289. 10.1016/j.cag.2020.07.010

Harrington, K. I., Zhao, Z., Schwartz, J., Kandel, S., Ermel, U., Paraan, M., Potter, C., & Carragher, B. (2024). Open-source Tools for CryoET Particle Picking Machine Learning Competitions (p. 2024.11.04.621608). bioRxiv. 10.1101/2024.11.04.621608

Hrabe, T., Chen, Y., Pfeffer, S., Kuhn Cuellar, L., Mangold, A.-V., & Förster, F. (2012). PyTom: A python-based toolbox for localization of macromolecules in cryo-electron tomograms and subtomogram analysis. Journal of Structural Biology, 178(2), 177–188. 10.1016/j.jsb.2011.12.003

Huang, Q., Zhou, Y., & Bartesaghi, A. (2024). MiLoPYP: Self-supervised molecular pattern mining and particle localization in situ. Nature Methods, 21(10), 1863–1872. 10.1038/s41592-024-02403-6

Jin, W., Zhou, Y., & Bartesaghi, A. (2024). Accurate size-based protein localization from cryo-ET tomograms. Journal of Structural Biology: X, 10, 100104. 10.1016/j.yjsbx.2024.100104

Khavnekar, S., Kelley, R., Waltz, F., Wietrzynski, W., Zhang, X., Obr, M., Tagiltsev, G., Beck, F., Wan, W., Briggs, J., Engel, B., Plitzko, J., & Kotecha, A. (2023). Towards the Visual Proteomics of C. reinhardtii using High-throughput Collaborative in situ Cryo-ET. Microscopy and Microanalysis, 29(Supplement_1), 961–963. 10.1093/micmic/ozad067.480

Kim, H. H.-S., Uddin, M. R., Xu, M., & Chang, Y.-W. (2023). Computational Methods Toward Unbiased Pattern Mining and Structure Determination in Cryo-Electron Tomography Data. Journal of Molecular Biology, 435(9), 168068. 10.1016/j.jmb.2023.168068

Lamm, L., Righetto, R. D., Wietrzynski, W., Pöge, M., Martinez-Sanchez, A., Peng, T., & Engel, B. D. (2022). MemBrain: A deep learning-aided pipeline for detection of membrane proteins in Cryo-electron tomograms. Computer Methods and Programs in Biomedicine, 224, 106990. 10.1016/j.cmpb.2022.106990

Lamm, L., Zufferey, S., Righetto, R. D., Wietrzynski, W., Yamauchi, K. A., Burt, A., Liu, Y., Zhang, H., Martinez-Sanchez, A., Ziegler, S., Isensee, F., Schnabel, J. A., Engel, B. D., & Peng, T. (2024). MemBrain v2: An end-to-end tool for the analysis of membranes in cryo-electron tomography (p. 2024.01.05.574336). bioRxiv. 10.1101/2024.01.05.574336

Last, M. G., Abendstein, L., Voortman, L. M., & Sharp, T. H. (2024). Streamlining segmentation of cryo-electron tomography datasets with Ais. eLife, 13, RP98552. 10.7554/eLife.98552

Liu, G., Niu, T., Qiu, M., Zhu, Y., Sun, F., & Yang, G. (2024). DeepETPicker: Fast and accurate 3D particle picking for cryo-electron tomography using weakly supervised deep learning. Nature Communications, 15(1), 2090. 10.1038/s41467-024-46041-0

Lucas, B. A., Himes, B. A., Xue, L., Grant, T., Mahamid, J., & Grigorieff, N. (2021). Locating macromolecular assemblies in cells by 2D template matching with cisTEM. eLife, 10, e68946. 10.7554/eLife.68946

Lučić, V., Rigort, A., & Baumeister, W. (2013). Cryo-electron tomography: The challenge of doing structural biology in situ. Journal of Cell Biology, 202(3), 407–419. 10.1083/jcb.201304193

Mahamid, J., Pfeffer, S., Schaffer, M., Villa, E., Danev, R., Kuhn Cuellar, L., Förster, F., Hyman, A. A., Plitzko, J. M., & Baumeister, W. (2016). Visualizing the molecular sociology at the HeLa cell nuclear periphery. Science, 351(6276), 969–972. 10.1126/science.aad8857

Majila, K., Arvindekar, S., Jindal, M., & Viswanath, S. (2025). Frontiers in integrative structural modeling of macromolecular assemblies. QRB Discovery, 6, e3. 10.1017/qrd.2024.15

Martinez-Sanchez, A. (2025). Template matching and machine learning for cryo-electron tomography. Current Opinion in Structural Biology, 93, 103058. 10.1016/j.sbi.2025.103058

Martinez-Sanchez, A., Kochovski, Z., Laugks, U., Meyer zum Alten Borgloh, J., Chakraborty, S., Pfeffer, S., Baumeister, W., & Lučić, V. (2020). Template-free detection and classification of membrane-bound complexes in cryo-electron tomograms. Nature Methods, 17(2), 209–216. 10.1038/s41592-019-0675-5

Maurer, V. J., Siggel, M., & Kosinski, J. (2024a). PyTME (Python Template Matching Engine): A fast, flexible, and multi-purpose template matching library for cryogenic electron microscopy data. SoftwareX, 25, 101636. 10.1016/j.softx.2024.101636

Maurer, V. J., Siggel, M., & Kosinski, J. (2024b). What shapes template-matching performance in cryogenic electron tomography in situ? Acta Crystallographica Section D: Structural Biology, 80(6), Article 6. 10.1107/S2059798324004303

Moebel, E., Martinez-Sanchez, A., Lamm, L., Righetto, R. D., Wietrzynski, W., Albert, S., Larivière, D., Fourmentin, E., Pfeffer, S., Ortiz, J., Baumeister, W., Peng, T., Engel, B. D., & Kervrann, C. (2021). Deep learning improves macromolecule identification in 3D cellular cryo-electron tomograms. Nature Methods, 18(11), Article 11. 10.1038/s41592-021-01275-4

Moreno, J. J., Martínez-Sánchez, A., Martínez, J. A., Garzón, E. M., & Fernández, J. J. (2018). TomoEED: Fast edge-enhancing denoising of tomographic volumes. Bioinformatics, 34(21), 3776–3778. 10.1093/bioinformatics/bty435

Ng, C. T., & Gan, L. (2020). Investigating eukaryotic cells with cryo-ET. Molecular Biology of the Cell, 31(2), 87–100. 10.1091/mbc.E18-05-0329

Peck, A., Yu, Y., Schwartz, J., Cheng, A., Ermel, U. H., Kandel, S., Kimanius, D., Montabana, E., Serwas, D., Siems, H., Wang, F., Zhao, Z., Zheng, S., Haury, M., Agard, D., Potter, C., Carragher, B., Harrington, K., & Paraan, M. (2024). Annotating CryoET Volumes: A Machine Learning Challenge (p. 2024.11.04.621686). bioRxiv. 10.1101/2024.11.04.621686

Pyle, E., & Zanetti, G. (2021). Current data processing strategies for cryo-electron tomography and subtomogram averaging. Biochemical Journal, 478(10), 1827–1845. 10.1042/BCJ20200715

Rice, G., Wagner, T., Stabrin, M., Sitsel, O., Prumbaum, D., & Raunser, S. (2023). TomoTwin: Generalized 3D localization of macromolecules in cryo-electron tomograms with structural data mining. Nature Methods, 20(6), Article 6. 10.1038/s41592-023-01878-z

Robinson, C. V., Sali, A., & Baumeister, W. (2007). The molecular sociology of the cell. Nature, 450(7172), 973–982. 10.1038/nature06523

Scheres, S. H. W. (2012). RELION: Implementation of a Bayesian approach to cryo-EM structure determination. Journal of Structural Biology, 180(3), 519–530. 10.1016/j.jsb.2012.09.006

Sofroniew, N., Lambert, T., Bokota, G., Nunez-Iglesias, J., Sobolewski, P., Sweet, A., Gaifas, L., Evans, K., Burt, A., Doncila Pop, D., Yamauchi, K., Weber Mendonça, M., Liu, L., Buckley, G., Vierdag, W.-M., Monko, T., Willing, C., Royer, L., Can Solak, A., … Zhao, R. (2025). napari: A multi-dimensional image viewer for Python (Version v0.6.4) [Computer software]. Zenodo. 10.5281/zenodo.16883660

Turk, M., & Baumeister, W. (2020). The promise and the challenges of cryo-electron tomography. FEBS Letters, 594(20), 3243–3261. 10.1002/1873-3468.13948

Uddin, M. R., Ahmed, A. Y., Tahmid, M. T., Alam, M. Z. U., Freyberg, Z., & Xu, M. (2024). TomoPicker: Annotation-Efficient Particle Picking in cryo-electron Tomograms (p. 2024.11.04.620735). bioRxiv. 10.1101/2024.11.04.620735

Uddin, M. R., Nguyen, T.-H., Tabib, H. M. S., Gandhi, K., & Xu, M. (2025). Unsupervised Multi-scale Segmentation of Cellular cryo-electron Tomograms with Stable Diffusion Foundation Model (p. 2025.06.25.661425). bioRxiv. 10.1101/2025.06.25.661425

Wagner, T., Merino, F., Stabrin, M., Moriya, T., Antoni, C., Apelbaum, A., Hagel, P., Sitsel, O., Raisch, T., Prumbaum, D., Quentin, D., Roderer, D., Tacke, S., Siebolds, B., Schubert, E., Shaikh, T. R., Lill, P., Gatsogiannis, C., & Raunser, S. (2019). SPHIRE-crYOLO is a fast and accurate fully automated particle picker for cryo-EM. Communications Biology, 2(1), Article 1. 10.1038/s42003-019-0437-z

Wagner, T., & Raunser, S. (2025). Cryo-electron tomography: Challenges and computational strategies for particle picking. Current Opinion in Structural Biology, 93, 103113. 10.1016/j.sbi.2025.103113

Wan, W., Khavnekar, S., & Wagner, J. (2024). STOPGAP: An open-source package for template matching, subtomogram alignment and classification. Acta Crystallographica Section D: Structural Biology, 80(5), Article 5. 10.1107/S205979832400295X

Xue, L., Lenz, S., Zimmermann-Kogadeeva, M., Tegunov, D., Cramer, P., Bork, P., Rappsilber, J., & Mahamid, J. (2022). Visualizing translation dynamics at atomic detail inside a bacterial cell. Nature, 610(7930), 205–211. 10.1038/s41586-022-05255-2

Zeng, X., Kahng, A., Xue, L., Mahamid, J., Chang, Y.-W., & Xu, M. (2023). High-throughput cryo-ET structural pattern mining by unsupervised deep iterative subtomogram clustering. Proceedings of the National Academy of Sciences, 120(15), e2213149120. 10.1073/pnas.2213149120

Zeng, X., Leung, M. R., Zeev-Ben-Mordehai, T., & Xu, M. (2018). A convolutional autoencoder approach for mining features in cellular electron cryo-tomograms and weakly supervised coarse segmentation. Journal of Structural Biology, 202(2), 150–160. 10.1016/j.jsb.2017.12.015

