## Supplementary material for "PickET: An unsupervised method for localizing macromolecules in cryo-electron tomograms"

### Supplementary Figures

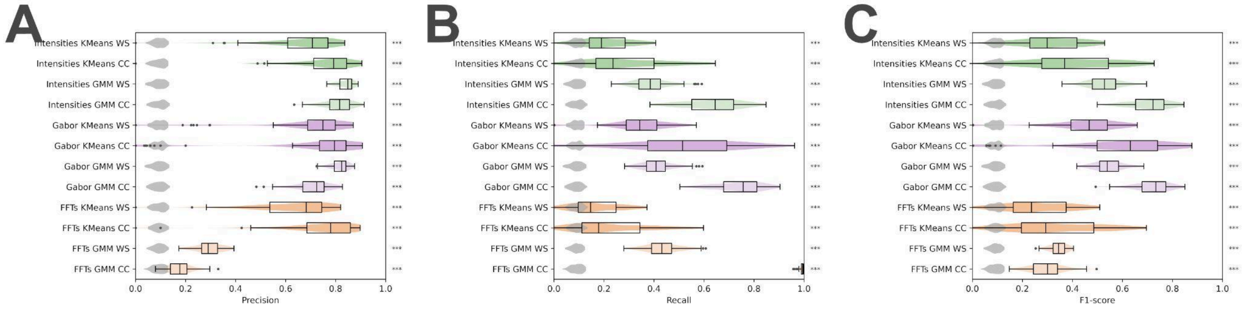


**Figure S1: Performance of the PickET library on simulated datasets.** (A) Precision, (B) recall, and (C) F1-score were computed on the particles localized by PickET and random predictions. The box and violin plots in panels A, B, and C compare different PickET workflows against a random baseline (grey violin plots) on a dataset comprising 88 simulated tomograms. The dots represent the outliers for the metrics measured on the model predictions. Green, purple, and orange violins represent the performance of the intensities, Gabor, and FFT workflows of PickET, respectively; darker (lighter) shades represent the KMeans (GMM) - based clustering workflows. The statistical significance of the difference between the metrics calculated on the model predictions and random predictions is represented by asterisks (‘n.s.’ - $p - value > 0.05$, ‘*’ - $0.05 \geq p - value > 0.01$, ‘**’ - $0.01 \geq p - value > 0.001$, and ‘***’ - $0.001 \geq p - value$). PickET predictions on 88 tomograms were used for these assessments.

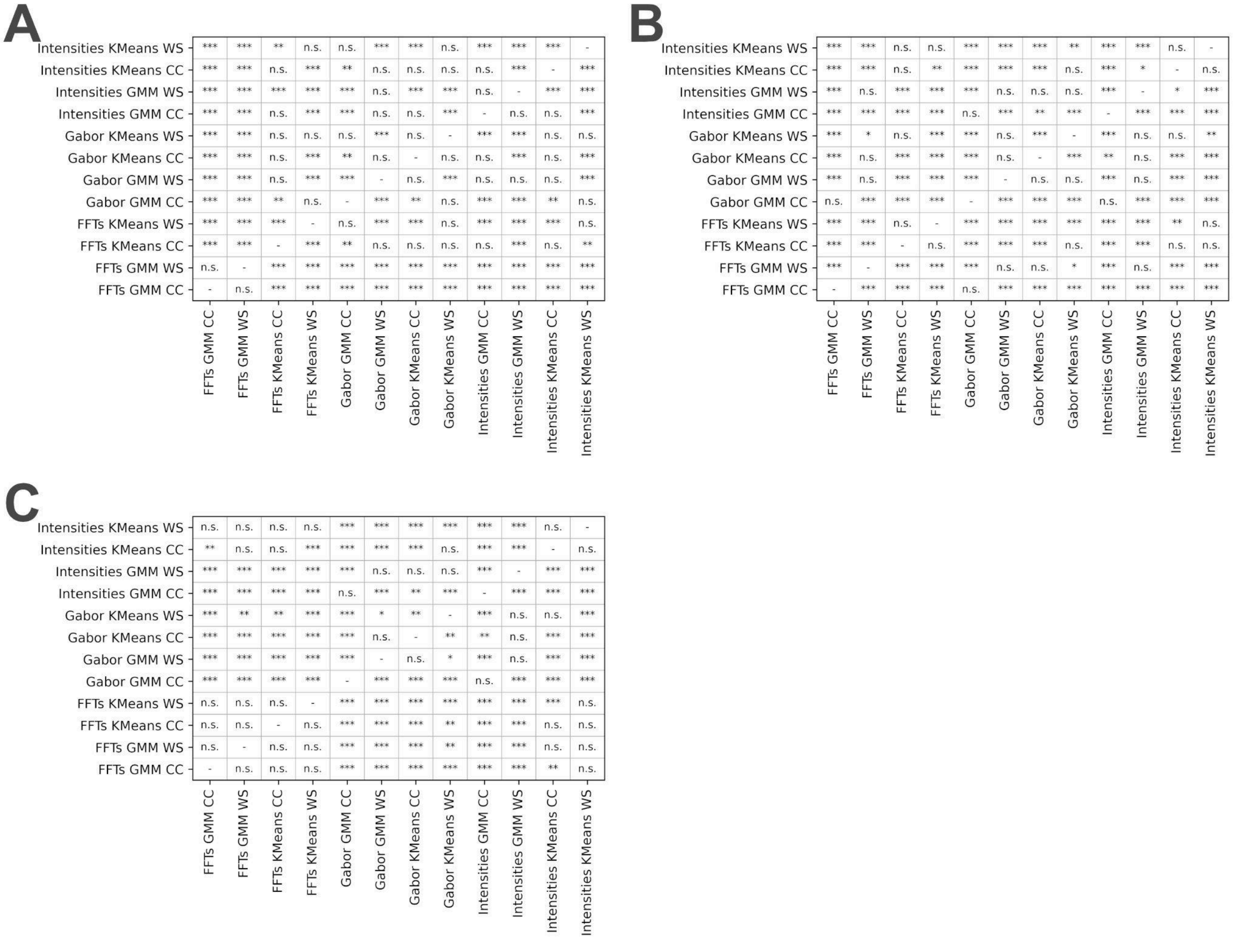


**Figure S2: Significance of the difference between the performance of PickET workflows on the simulated dataset.** The statistical significance of the difference in the precision (A), recall (B), and F1-score (C) between pairs of PickET workflows is shown in terms of Bonferroni-adjusted p-values obtained from a post hoc Dunn's test performed following a significant Kruskal-Wallis test (p-value < 0.05). The statistical significance of the difference between the distributions of the metrics calculated on the predictions from two different workflows is represented by asterisks ('n.s.' - $p - value > 0.05$, '*' - $0.05 \geq p - value > 0.01$, '**' - $0.01 \geq p - value > 0.001$, and '***' - $0.001 \geq p - value$). Self comparisons are marked with '-'. PickET predictions on 88 tomograms were used for these assessments.

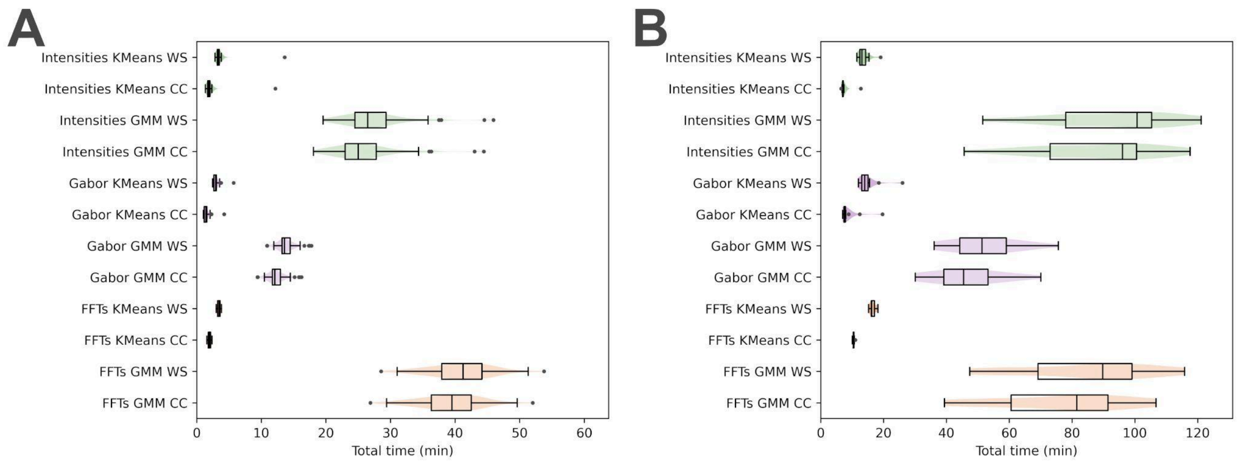


**Figure S3: Efficiency of the PickET library on a simulated and a real-world dataset.** (A) shows the time taken by each of the PickET workflows on eight simulated $512 * 512 * 200$ voxel tomograms on a workstation running an AMD Ryzen Threadripper 3990X 64-Core Processor with 2.9 GHz clock speed, along with 264 GB RAM and an NVIDIA RTX 6000 Ada Generation GPU. (B) shows the time taken by each of the PickET workflows on ten real-world $1024 * 1024 * 512$ voxel tomograms from the CZI-DS-10301 on a cluster node running an AMD Epyc 7763 64-Core Processor with 1.5 GHz clock speed, along with 528 GB RAM and an NVIDIA RTX A6000 GPU.

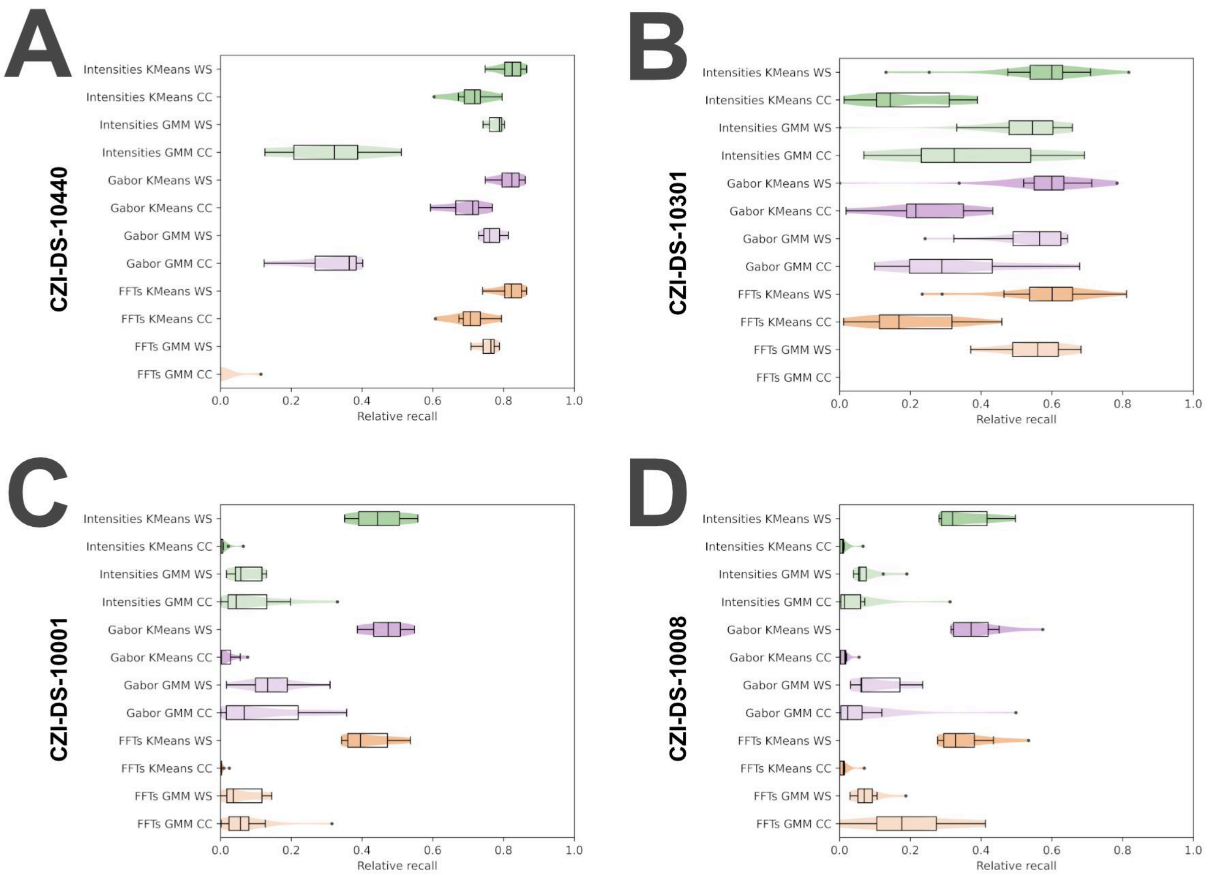


**Figure S4: Relative recall on PickET predictions on real-world datasets.** Relative recall was computed on the particle localizations predicted by PickET on (A) CZI-DS-10440 $(n = 7)$, (B) CZI-DS-10301 $(n = 18)$ (for FFTs-GMM-CC $(n = 6)$), (C) CZI-DS-10001 $(n = 10)$, and (D) CZI-DS-10008 $(n = 9)$ datasets. The dots represent the outliers for the metrics measured on the model predictions. Green, purple, and orange violins represent the performance of the intensities, Gabor, and FFT workflows of PickET, respectively; darker (lighter) shades represent the KMeans (GMM) - based clustering workflows. The statistical significance of the difference between the metrics calculated on the model predictions and random predictions is represented by asterisks ('n.s.' - $p - value > 0.05$, '*' - $0.05 \geq p - value > 0.01$, '**' - $0.01 \geq p - value > 0.001$, and '***' - $0.001 \geq p - value$).

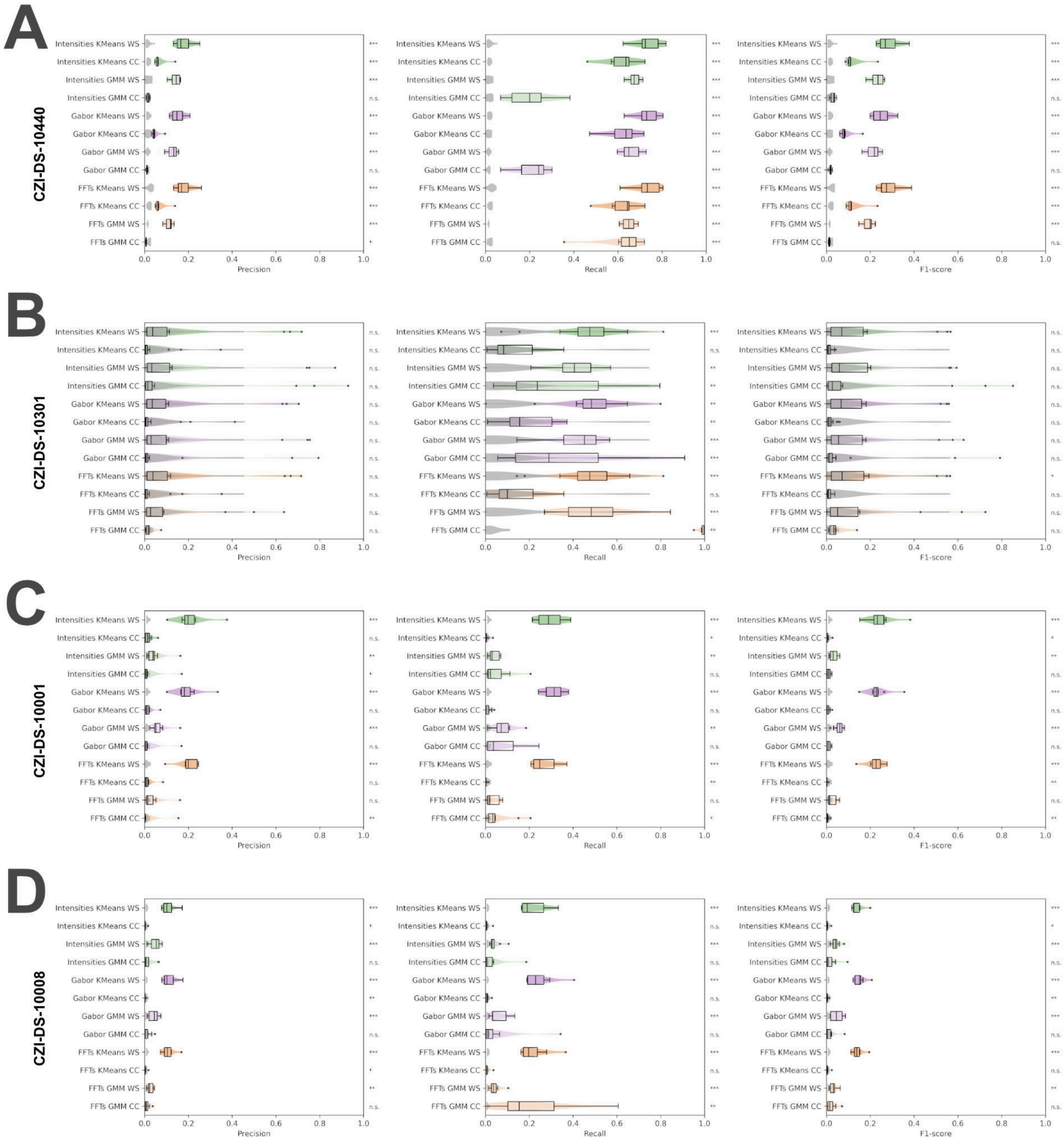


**Figure S5: Precision, Recall, and F1-score of PickET on real-world datasets.** Precision (left), recall (center), and F1-score (right) were computed on the particle localizations predicted by PickET on (A) CZI-DS-10440 ($n = 7$), (B) CZI-DS-10301 ($n = 18$) (for FFTs-GMM-CC ($n = 6$)), (C) CZI-DS-10001 ($n = 10$), and (D) CZI-DS-10008 ($n = 9$) datasets. The dots represent the outliers for the metrics measured on the model predictions. Green, purple, and orange violins represent the performance of the intensities, Gabor, and FFT workflows of PickET, respectively; darker (lighter) shades represent the KMeans (GMM) - based clustering workflows. The statistical significance of the difference between the metrics calculated on the model predictions and random predictions is represented by asterisks ('n.s.' - $p - value > 0.05$, '*' - $0.05 \geq p - value > 0.01$, '**' - $0.01 \geq p - value > 0.001$, and '***' - $0.001 \geq p - value$).

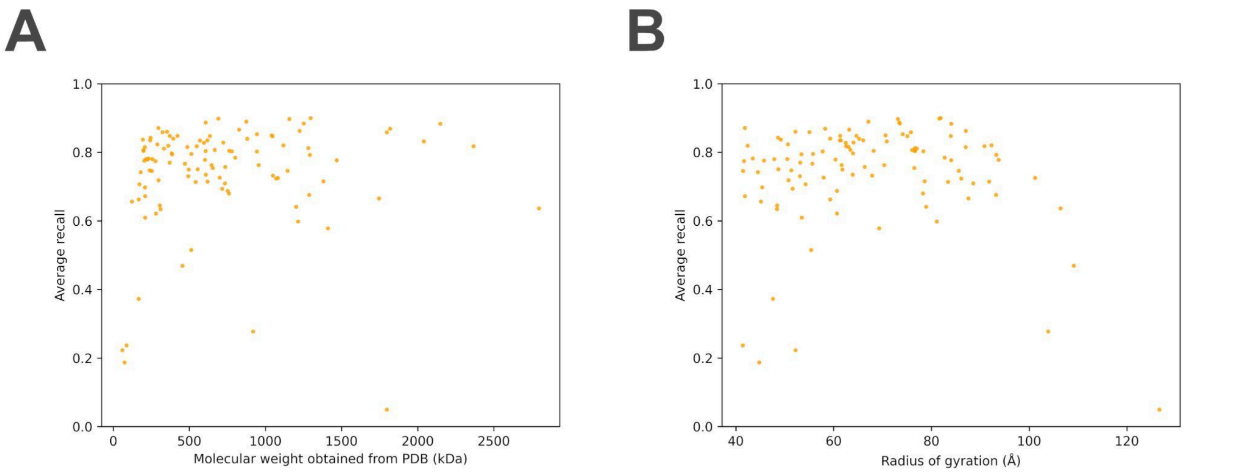


**Figure S6: Dependence of PickET performance on particle characteristics.** The average particle-wise recall on the predictions from PickET is shown against the molecular weight (A) and radius of gyration (B) of the macromolecules.

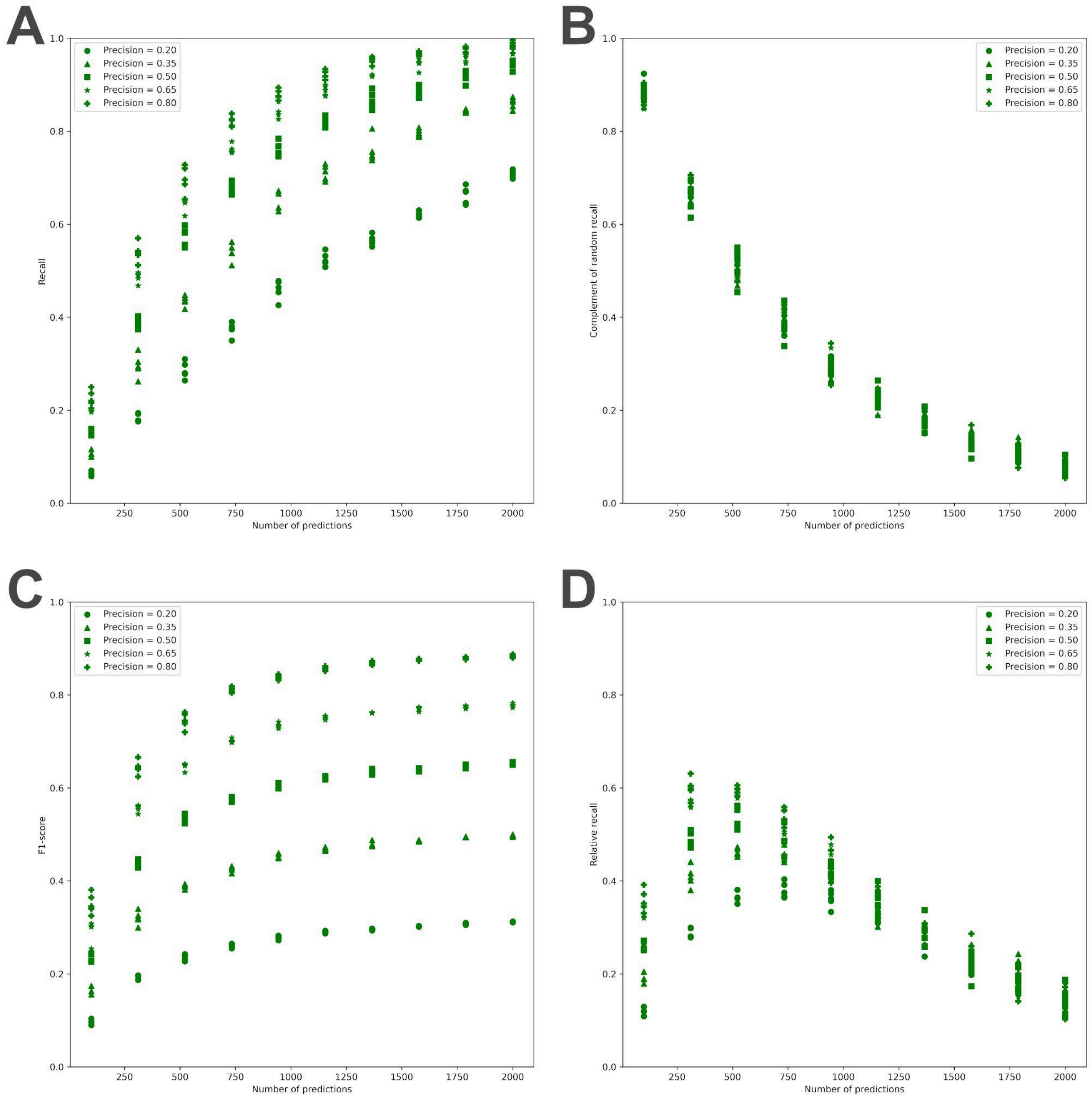


**Figure S7: Comparison of relative recall and F1 score on a simulated dataset.** The particle localization task was simulated on a sample dataset comprising 500 randomly initialized ground truth particles in a square of length 100 pixels. A model was devised to pick a variable number of particles from this square while keeping the precision fixed. (A) depicts the relationship between recall and the number of predicted particles, (B) depicts the relationship between complement of random recall (See **Materials and Methods, Assessments**) and the number of predicted particles, (C) depicts the relationship between the F1-score and the number of predicted particles, and (D) depicts the relationship between the relative recall and the number of predicted particles. The different markers (circle, triangle, square, star, and plus) in the scatter plots represent uniformly spaced values of precision between 0.2 and 0.8.

### Supplementary Table

| Dataset | Macromolecule | Recall |
|---|---|---|
| CZI-DS-10001 | Cytosolic ribosome | 0.31 |
| | Fatty acid synthase | 0.20 |
| CZI-DS-10008 | Cytosolic ribosome | 0.27 |
| | Hydrogen-dependent $CO_2$ reductase filament | 0.09 |
| CZI-DS-10301 | Cytosolic ribosome | 0.54 |
| | Microtubule | 0.56 |
| | Mitochondrial proton-transporting ATP synthase | 0.50 |
| | Nucleosome | 0.46 |
| | Ribulose bisphosphate carboxylase | 0.46 |
| CZI-DS-10440 | Beta amylase | 0.65 |
| | Beta galactosidase | 0.83 |
| | Cytosolic ribosome | 0.81 |
| | Ferritin | 0.56 |
| | PP7-VLP | 0.83 |
| | Thyroglobulin | 0.83 |

**Table S1: Average particle-wise recall of PickET on real-world datasets.**
